# Immunologic comparisons of strain and induction method in an improved mouse model of intrauterine fibrosis

**DOI:** 10.1101/2024.10.30.621047

**Authors:** Jamie L. Hernandez, Jonathan Daniel, Jessica L. Stelzel, Neeti R. Prasad, Vance V. Soares, Joshua C. Doloff

## Abstract

Intrauterine adhesions are growths of fibrotic tissue within the uterine cavity and can arise from a variety of tissue-damaging stimuli. Immune cells are known to mediate fibrotic responses, but specific mechanisms require further elucidation. Here, we compared intrauterine fibrosis development and immune responses across different mouse strains and induction methods. We aimed to identify a consistent and more clinically relevant mouse model of intrauterine fibrosis, whether immune responses differ in response to different stimuli, and which potential key immune cell populations are responsible for intrauterine fibrosis susceptibility. Intrauterine fibrosis induction methods were compared using surgical curettage or transcervically administered chemical (quinacrine) models. Measurements of tissue morphology and collagen gene expression indicate BALB/c mice are more susceptible than C57BL/6 mice to intrauterine fibrosis. In chemically induced BALB/c uterine tissues, gene expression and flow cytometry data show greater pro-inflammatory macrophage responses, implicating a possible role in fibrogenesis consistent with human intrauterine adhesion data. Findings from this study demonstrate the importance of mouse strain selection in studies of intrauterine adhesions. Furthermore, we show that a new hormone-synchronized, chemically induced mouse model can more uniformly and reliably provoke fibrotic tissue response. This model may allow for greater elucidation of mechanisms involved in intrauterine adhesion development, and exploratory therapeutic studies for treatment intervention.

## 1. Introduction

Intrauterine adhesions (IUA), also known as Asherman’s syndrome when severe, is the growth of fibrotic tissue within the uterine cavity. Patients with IUA can experience amenorrhea, recurrent pregnancy loss, and infertility.^1,2^ Current treatment options include surgical removal of fibrotic tissue, but IUA regrowth is common.^1^ Although often referred to as a rare disease,^3^ actual occurrence rate is relatively unknown and understudied due to variations in symptom presentation.^1^ IUA arises from tissue damage of the endometrium, occurring in up to 4% of patients after cesarian section, 4% of patients undergoing postpartum dilation and curettage (D&C), and 7% of patients undergoing D&C for early spontaneous abortion.^1^ Endometrial tissue damage and resulting intrauterine fibrosis is also known to occur from infections^4^ and from chemical sclerosing agents studied for non-surgical female permanent contraception.^5–9^ Fibrotic diseases like IUA develop from excess extracellular matrix production, especially collagens, by activated fibroblasts.^10–12^ Immune cells are known to respond to tissue damage and mediate fibroblast behavior towards either pro-fibrotic or pro-wound healing reactions.^11,12^ However, the specific mechanisms and key immune populations involved in intrauterine fibrosis are not fully understood.^2^ Improved models of IUA and intrauterine fibrosis more generally are needed to improve understating of immune mechanisms and identify treatment targets.

Although measurements of human biopsies are promising for clinical relevance, animal models are necessary for gaining greater insight into pathologies like IUA, especially, in many cases, due to limited availability of patient samples. Moreover, mouse models have a wide variety of tools available to study causal relationships of their biology, allow for the study of early tissue responses, and enable screening experiments of promising treatments. IUA has been studied in mice using a curettage model, where the lumen of the uterine horn is surgically damaged using a needle.^2,13–17^ Cyclic changes of female sex hormones, like estrogens and progesterone, are known to affect uterine tissue morphology and local immune cell populations.^18^ Therefore, rodent models of uterine fibrosis often monitor estrous cycle stage daily, and control for induction at diestrus,^17,19,20^ when relative serum concentrations of estradiol is low and progesterone is intermediate.^21^ However, other studies do not control for cyclicity.^13–16^ Another inconsistency in IUA mouse models is the strain used.^13,14,16,17^ Differences in genetic background are known to cause organ-specific susceptibilities to fibrosis induction in other fibrotic disease models, but the effect of strain on intrauterine fibrosis susceptibility is unknown.^22^ These genetic differences cause biases in immune reactions, with C57BL/6 mice generally having more pro-inflammatory Th1-like responses, while BALB/c mice have anti-inflammatory or pro-tissue growth Th2-like responses.^22,23^ Understanding the effect of mouse strain on intrauterine fibrosis therefore is necessary to identify the most appropriate model to use in mirroring clinically observed pathologies, and would provide greater insight into the immune mechanisms that lead to fibrosis susceptibility or resistance. Finally, mouse models of other fibrotic tissue diseases often employ chemical agents to induce fibrosis.^22^ Assessments of chemical methods in intrauterine fibrosis therefore could present a new model for studying intrauterine fibrosis, and comparisons to curettage models could inform whether observed immune mechanisms are common based on tissue or whether they arise under different induction method-specific pathways.

In this study, we investigated susceptibility and immune mechanisms of intrauterine fibrosis using curettage and chemically induced methods, and across C57BL/6 and BALB/c mice. We hypothesized that there would be differences in the magnitude of fibrosis induction between strains. In characterizing the fibrotic responses from these models, we also aimed to describe the differences or commonalities of immune responses by gene expression analyses and by flow cytometry. Findings from this research can inform the most appropriate use of mouse strain for future IUA and intrauterine fibrosis research. Moreover, we present a simple and effective alternative, an estrus-synchronized and chemically induced mouse model, to enable more studies of this pathology. Finally, characterization of observed immune response may also suggest potential therapeutic drug targets for further study.

## 2. Results and Discussion

### 2.1. Inconsistencies in response are seen in non-hormone-controlled induction method screening studies

Initial studies were run to screen standard and new chemical methods for intrauterine fibrosis induction in mice. Mouse estrous cyclicity was not controlled for, and only C57BL/6 mice were used as this strain is generally more pro-fibrotic for other disease models^22^ and has been used in previous IUA studies.^14^ Curettage induction is the standard method used to study IUA in mice, and mimics clinical procedures by mechanically damaging the tubal lumen with a needle. Chemical induction of intrauterine fibrosis has been used clinically and in other animal models; however, the small scale and anatomy of the mouse female reproductive tract has been limiting for such studies in mice (**Fig. 1A**). Silver nitrate is one studied sclerosing agent which has shown potency in non-human primate and guinea pig models.^6,24^ Quinacrine is the most studied uterine-sclerosing agent used in clinical permanent contraception studies; however current clinical use is limited due to debated concerns of safety.^7,20,25^ To overcome the earlier mentioned limitations of small mouse anatomy and deliver these agents to the uterine horns, we used a mouse-scale transfer device designed for artificial insemination studies (**Fig. 1B**).^26^ Unlike the curettage model, chemical induction procedures with the transcervical transfer device were non-surgical. Initial tests showed that a single transcervical administration could deliver solution to both uterine horns (**Fig. 1C**). High and low dosages of the sclerosing agents were determined from previous animal studies with allometric scaling calculations.^6,7,20,24,27^ Previous studies of quinacrine also showed greater tissue changes after two administrations of the agent,^20,28^ and since multiple administrations were possible with transcervical delivery, quinacrine was dosed at the beginning of the study and again after 14 days (**Fig. 1D**).

**Figure 1.**
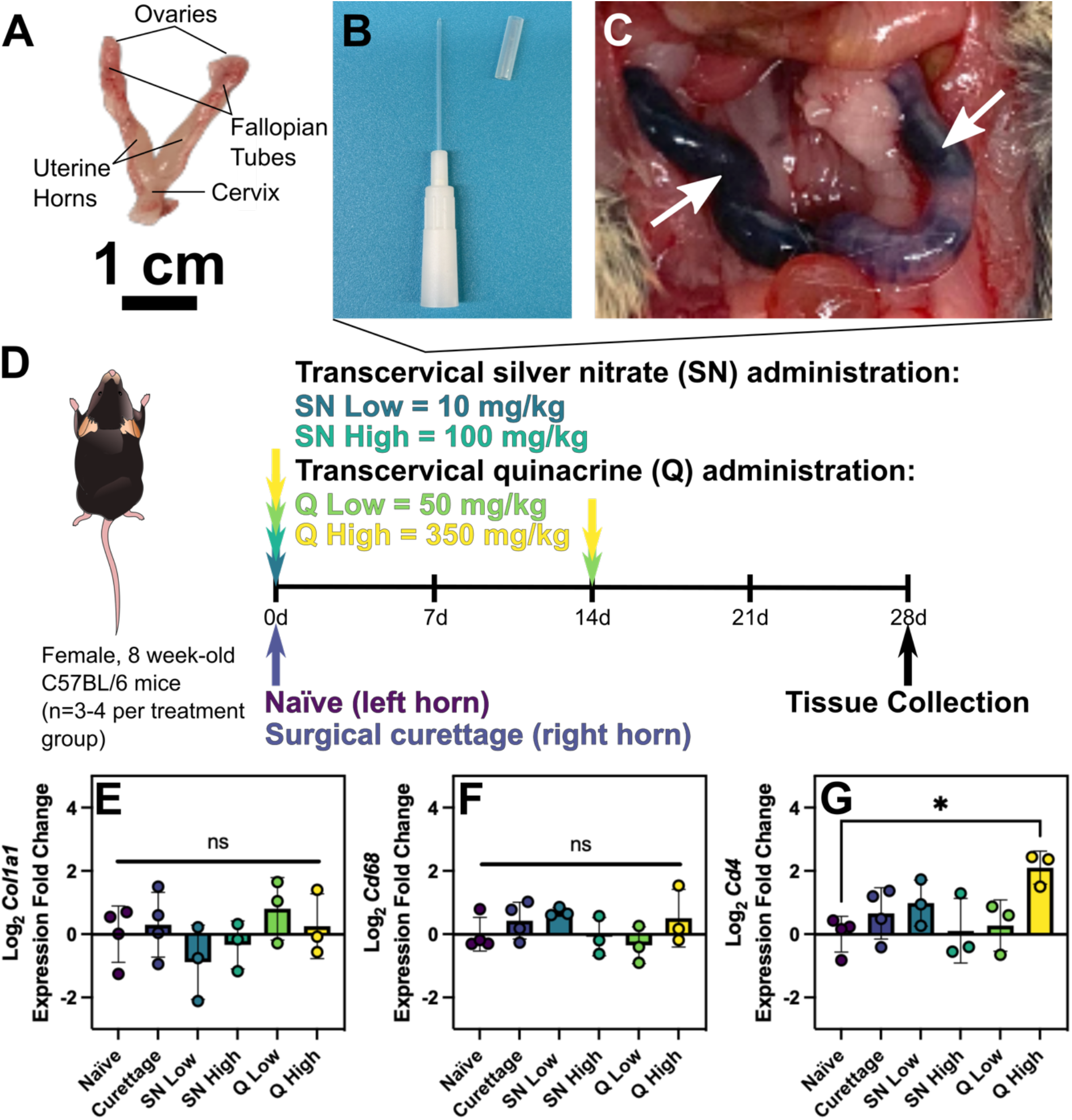
Hormone synchronization is necessary for consistent fibrotic responses. Chemical sclerosing agents were administered (A) to the mouse female reproductive tract using (B) a mouse scale commercial transfer device (ParaTech C&I device). (C) Transcervical administration of Trypan Blue shows distribution of solutions (white arrows) across both uterine horns with a single administration. (D) Without cycle synchronization,8-week-old female C57BL/6 mice underwent surgical curettage of the right uterine horn, or low/high dosing of the sclerosing agents silver nitrate (SN) or quinacrine (Q). Uterine horn tissues were assessed for relative gene expression of (E) Col1a1, (F) Cd68, and (G) Cd4 (n=3-4 mice, with n=1 uterine horn sections averaged for naïve and curettage mice or n=2 uterine horn tissues averaged for transcervically delivered sclerosing agents, plotted as the Log2 transform normalized to naïve uterine horn tissues (plotted, mean ± standard deviation as a measure of dataset variability)).Statistical significance determined by ordinary one-way ANOVA with Tukey’s multiple comparisons test, where ns=p>0.05 and *=p<0.05.

All induction methods and dosages were well tolerated. Some indications of fibrosis induction were observed via histology for high-dose quinacrine treatments (data not shown); however, the response was inconsistent across replicates. Gene expression for matrix proteins (collagen 1a1, *Col1a1*, **Fig. 1E**), macrophages (*Cd68*, **Fig. 1F**), and CD4 helper T cells (*Cd4*, **Fig. 1G**) was measured by qPCR to further assess fibrotic responses in uterine tissues. Without controlling for sex hormones and cyclicity, responses were inconsistent, with high standard deviation in relative gene expression and non-significant differences in expression levels relative to naïve control tissues. This is with the exception of elevated *Cd4* expression in high-dose, quinacrine-induced tissues relative to naïve (p=0.0253). These data warrant further studies of induction models, especially the standard curettage model and transcervically delivered quinacrine, but we hypothesized that control over cyclicity was necessary to improve consistency.

### 2.2. Curettage and chemical induction methods are effectively administered with DMPA estrous cycle synchronization treatments

Fluctuations of female sex hormones due to the menstrual cycle alter immune cell populations;^18^ therefore, timing may affect fibrosis susceptibility or resistance for our induction model. Mouse estrous cycles differ from human menstrual cycles, having approximately four-day instead of 28-day cycles (**Fig. 2A**).^21^ Additionally, the increased frequency of the mouse estrous cycle means there is greater variation of hormone and immune cell levels throughout the typical four-to-six-week wound healing, or fibrogenesis, timeframe.^29^ Administration of the long-acting, synthetic progesterone depot medroxyprogesterone (DMPA) is known to control cyclicity of female sex hormones in humans and mice, having initial increases of serum progestogen concentrations after administration, which slowly decrease over time for physiological concentrations beyond 28-days (**Fig. 2A**).^30^ We hypothesized that administration of DMPA prior to induction experiments would improve induction consistency and translatability to humans.

**Figure 2.**
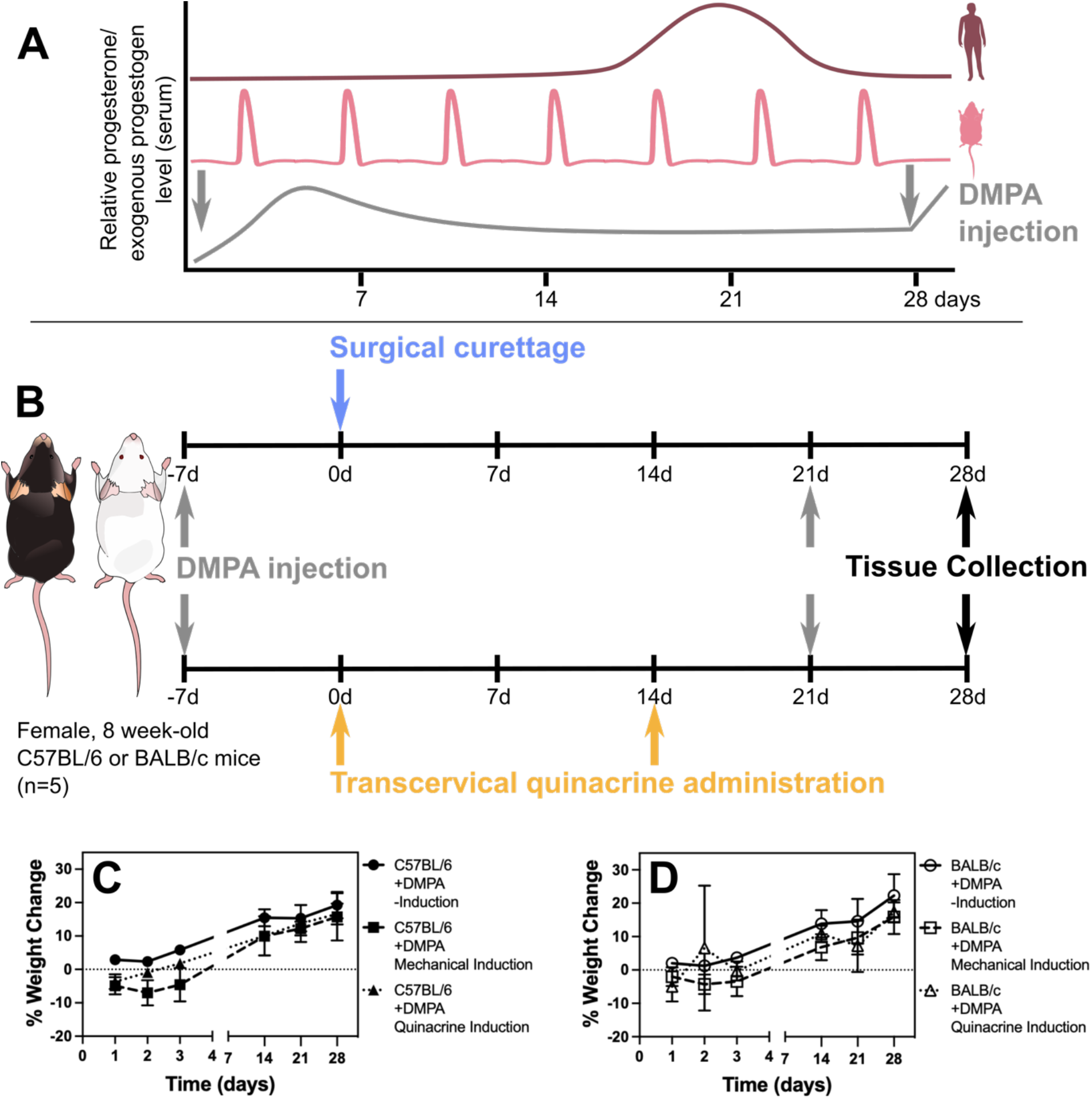
DMPA cycle synchronization assessed in surgical curettage and transcervical quinacrine induction models in both C57BL/6 and BALB/c mice. Relative blood serum hormone concentrations vary cyclically over time as shown in (**A**) a schematic of progesterone/progestogen levels for humans, mice, and either species under DMPA treatments over 28-days.^21,30^ (**B**) Intrauterine fibrosis was induced in female, 8-week-old C57BL/6 or BALB/c mice using one-time surgical curettage to both left and right uterine horns or twice administered quinacrine, with DMPA injections administered to all mice 7-days prior to induction and then every 28 days. Trends in % weight change following induction methods are measured for (**C**) C57BL/6 and (**D**) BALB/c mice (n=5 mice per treatment group, plotted for mean ± standard deviation to show dataset variation).

Due to the known effect of mouse strain on other fibrotic diseases, and the unknown effect on intrauterine fibrosis, we assessed the standard curettage model and the promising quinacrine chemical induction method in both C57BL/6 and BALB/c mice with DMPA synchronization treatments (**Fig. 2B**). To generate control specimens for comparison, mice from both strains underwent DMPA treatments but no induction procedure. Surgical and transcervical procedures were well tolerated in all mice, as demonstrated by body score and weight, both of which were stable and even increased as animals aged over time (**Fig. 2C&D**). There were no statistical differences in weight change between C57BL/6 and BALB/c mice for all comparison groups (control, p=0.7045; curettage, p=0.5689; and quinacrine, p=0.6801). Both induction methods caused initial decreases in weight change, most significantly different for curettage, in C57BL/6 mice. Further, while lesser effects were seen in BALB/c mice, curettage induction yielded significantly decreased weight change after two days instead of one (all p-values shown in **Supplemental Table 1**). Overall, DMPA administration proved to be a relatively simple and well tolerated method for controlling cyclicity prior to induction.

**Table 1.**
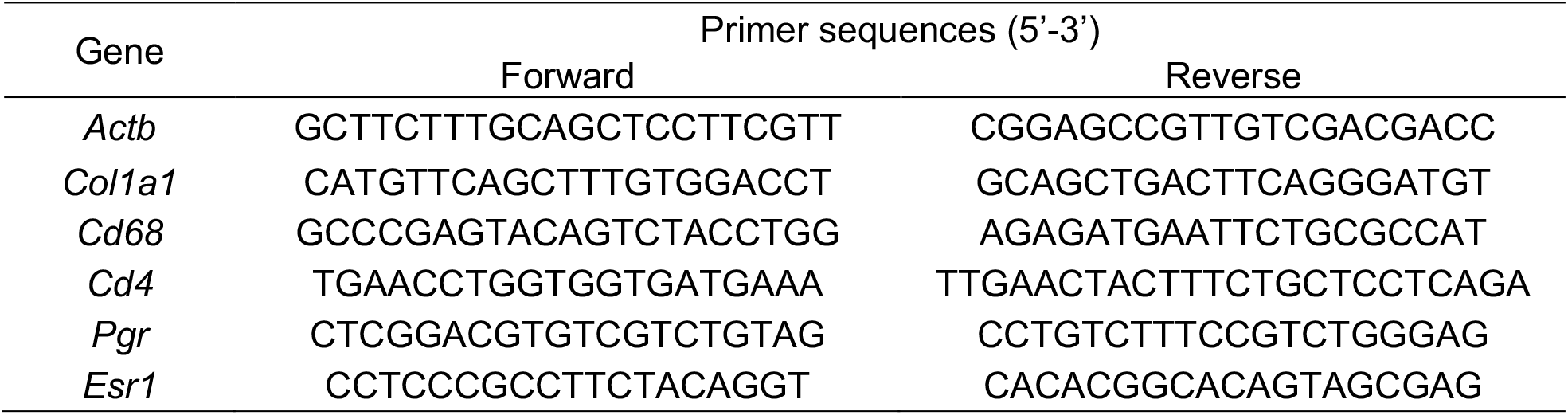
Primers and sequences used for qPCR.

### 2.3. Anatomical and histological changes are observed after induction, especially for quinacrine-treated BALB/c mice

Gross anatomical tissue changes following induction treatments could be seen 28 days after induction procedures (**Fig. 3A-F**). Specifically, regions of edema and accumulation of intrauterine luminal fluids were observed in all induction treatment groups. Hydrosalpinx, or distension of the fallopian tubes from fluid accumulation, can arise in humans from scar tissue occlusion of the tubal lumen, commonly caused by pelvic inflammatory disease.^31^ Our observing regions of edema (**Fig. 3B, C, & E**) may confirm induction of inflammatory tissue damage responses, and considering the small diameter of the mouse uterine horn, may also indicate possible fibrotic occlusion of the tubal lumen.

**Figure 3.**
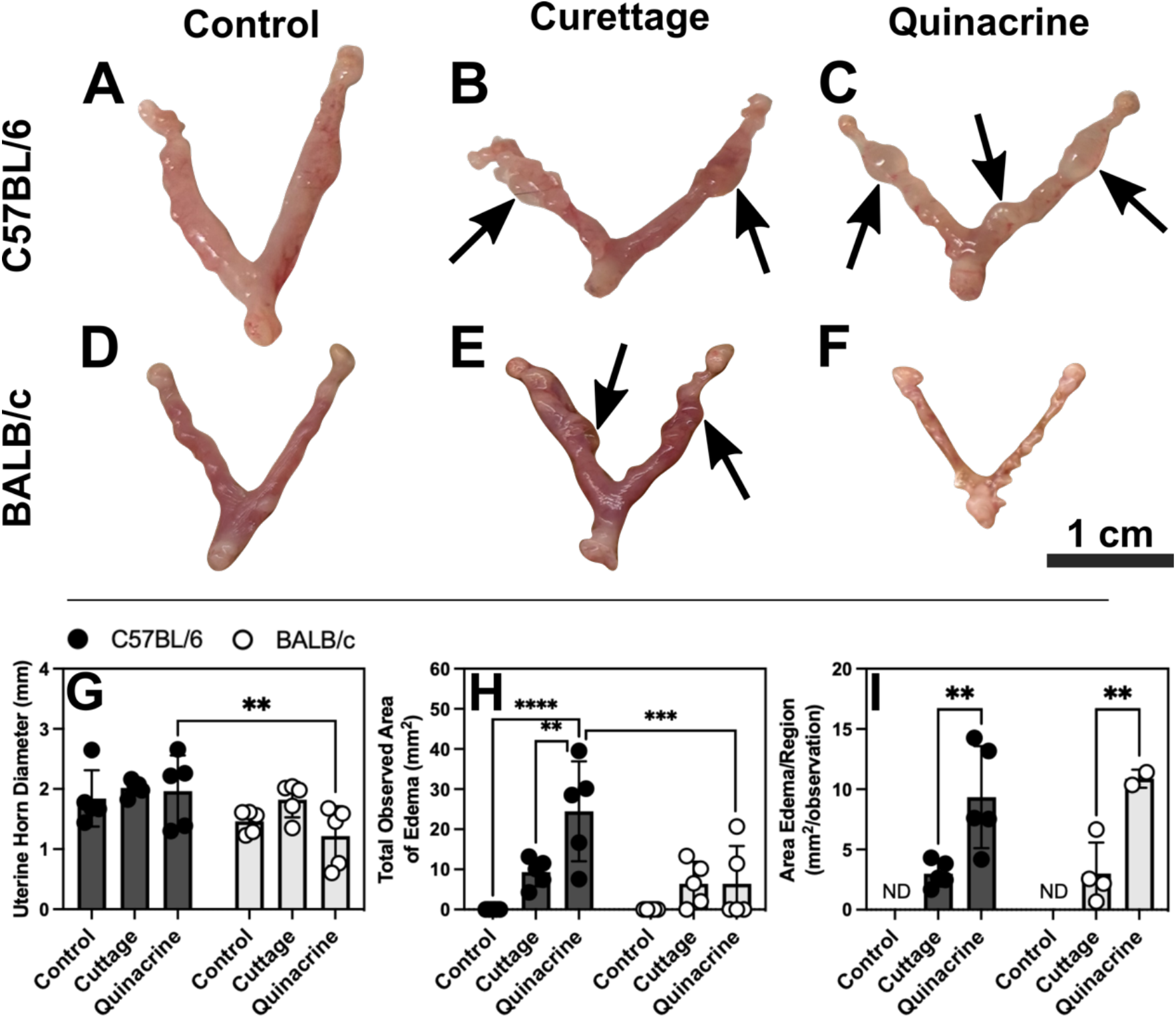
Induction methods induce intrauterine tissue changes at 28 days. Representative female reproductive tract images for (**A-C**) C57BL/6 and (**D-F**) BALB/c mice (**A&D**) without induction, (**B&E**) with surgical curettage, or (**C&F**) with transcervical quinacrine induction treatment are shown, with black arrows marking observed regions of edema. Images are captured via digital camera with scale bars representing 1 cm. Image analyses of uterine horn images quantified (**G**) uterine horn diameter, (**H**) total observed area of edema, and (**I**) area of edema per observed region of edema are plotted as average ± standard deviation to show variation in measured images within the sample group (n=5 mouse female reproductive tracts, with n=6 averaged measurements for diameter). Statistical analysis conducted by ordinary two-way ANOVA and Tukey’s Multiple comparisons test, or Fisher’s LSD test for analyses which have groups of edema not detected (ND), with **=p<0.01, ***=p<0.001, and ****=p<0.0001.

Curettage-induced mice consistently had regions of edema located near the superior region of the uterine horns, where the needle was inserted into the lumen to induce damage (**Fig. 3B&E**). Chemically (quinacrine) induced tissues had larger regions of accumulated luminal fluids (**Fig. 3C**). In quinacrine-treated BALB/c mice, three of five female reproductive tracts did not have observed edema, but instead appeared thinner and discolored suggesting a different tissue response not observed for the other groups (**Fig. 3F**). Image analysis measurements of tubal diameter capture the overall significant decrease for quinacrine-treated BALB/c tissues compared to C57BL/6 (p=0.0069, **Fig. 3G**). Moreover, total measured area of edema was greatest for quinacrine-treated C57BL/6 mice, and even significantly greater than quinacrine-treated BALB/c mice (p=0.0004, **Fig. 3H**). When regions of edema were detected, the area per region was greatest in both strains for quinacrine treatments compared to curettage (p=0.004 for C57BL/6, p=0.0076 for BALB/c, **Fig. 3I**). Assessed induction models appear to induce fibrotic changes, which warranted further quantification.

Histological evidence of uterine fibrosis development includes disruption of the epithelium,^13,15^ tissue occlusion or adhesion of the tubal lumen,^19,20,24^ and/or growth of fibrotic masses.^17^ Uterine horn cross-sections show possible evidence of these tissue changes for induction treatment mice (**Fig. 4A-F**). Due to the natively collagen rich extracellular matrix of uterine tissues, H&E stained sections can better illustrate tissue changes, and were therefore used for image insets. Control histology show open tubal lumens with intact epithelium (**Fig. 4A&D**). Although the uterine tissue is natively collagen rich, stained blue in Masson’s Trichrome images, fibrotic masses can be observed within the tubal lumen in some histology sections of C57BL/6 curettage mice (**Fig. 4B**). For chemically induced mice, possible evidence of epithelium disruption and cellular infiltrate within the tubal lumen are seen in C57BL/6 mice (**Fig. 4C**). Furthermore, possible evidence of tubal occlusion is seen for chemically induced BALB/c mice, consistent with gross anatomical observations (**Fig. 4F**). Specifically, uterine horn histology cross-sections were observed to be smaller for BALB/c chemically induced tissues, consistent with the smaller uterine horn diameter measurements shown in **Fig 3G**. The constricted diameter of the uterine horn tissue is hypothesized to be another manifestation of a fibrotic response. Since histology images are location-dependent, holistic measurements of tissue responses are necessary to capture overall uterine responses.

**Figure 4.**
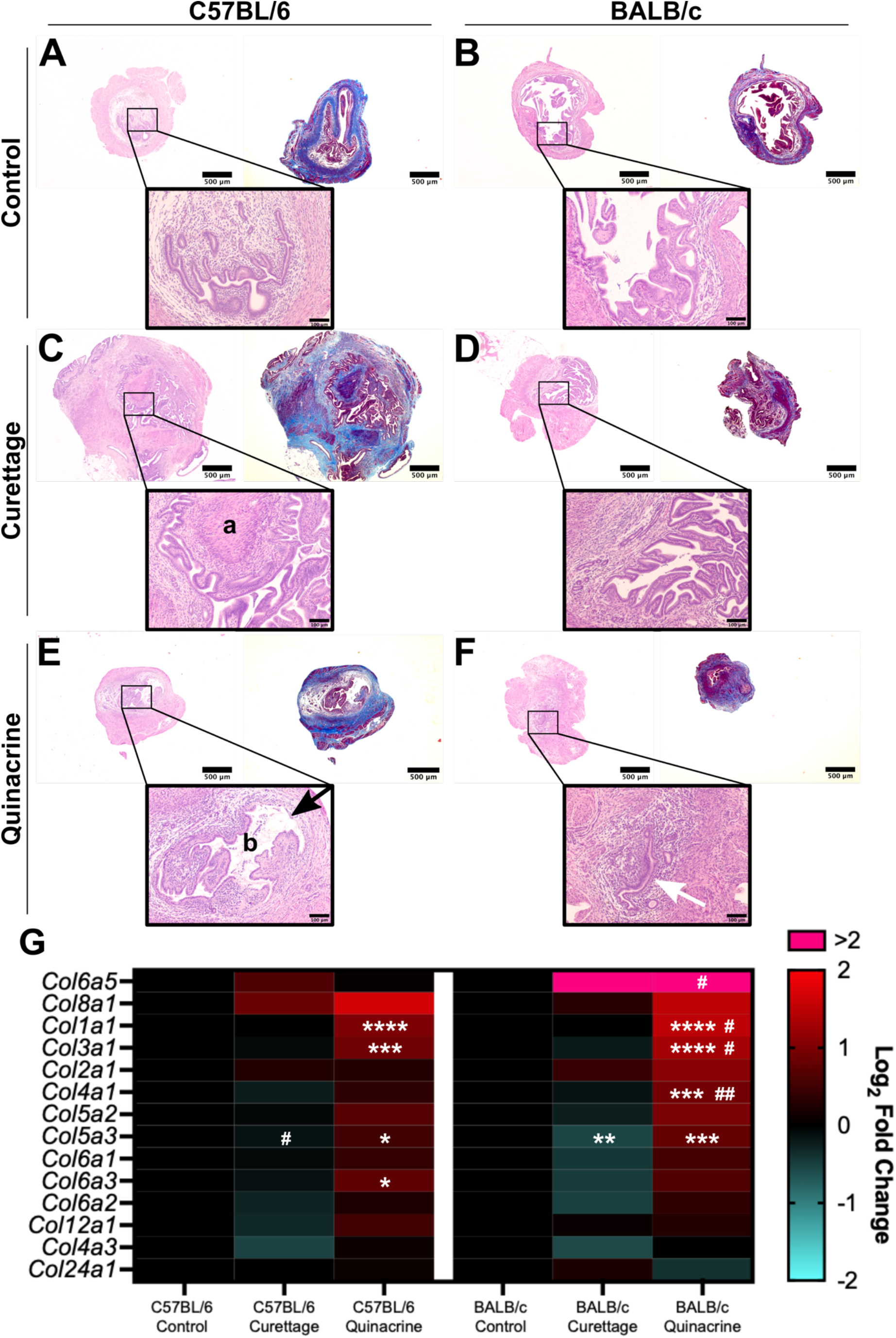
Quinacrine induction methods lead to increased collagen gene expression and histological changes to uterine horn lumen. Representative histology images of uterine horn cross-sections are shown for DMPA-treated (**A, C, & E**) C57BL/6 and (**B, D, & F**) BALB/c mice (**A&B**) without induction, (**C&D**) with surgical curettage, or (**E&F**) with transcervical quinacrine induction treatment. Images are captured by brightfield microscopy, and are shown at 4x magnification for H&E (left), Masson’s Trichrome (right), and H&E at 20x magnification (inset). Scale bars, 500 μm. Regions of possible epithelium disruption (black arrow), occlusion of the lumen (white arrow), fibrotic growth (a), and cellular infiltrate (b) are marked. Tissue matrix modifications are also assessed by (**G**) collagen gene expression quantified by NanoString and represented as average Log2 fold change normalized to the strain specific control (DMPA treated/no induction) (n=4 mice). Statistical significance was determined by ordinary two-way ANOVA with Tukey’s multiple comparisons test. Significant differences in gene expression compared to strain-specific control group are shown by *=p<0.05, ***=p<0.001, and ****=p<0.0001. Significant increases in gene expression compared across mouse strain for the same treatment are shown by ^#^=p<0.05 and ^##^=p<0.01.

To quantify fibrotic responses, we measured the expression of numerous collagen isoforms as part of a custom-built multiplexed NanoString gene expression panel (**Fig. 4G**). All measured genes can be seen in **Supplemental Fig. 1**. Chemically induced mice, especially BALB/c, show the greatest and most significant expression of collagen genes. Expression of basement membrane associated and epithelial cell secreted type VI collagen^32^ gene *Col6a5* shows the largest average relative magnitude of the assessed collagen genes in induced BALB/c mice (Log2 fold change = 2.26x and 3.52x for curettage and quinacrine, respectively), with significantly higher expression for quinacrine-treated BALB/c mice than C57BL/6 mice (p=0.0239, all p-values shown in **Supplemental Table 2**). However, the response is not consistent across biological replicates, and not significantly greater than control (no induction, DMPA-treated) mice. Type I and III collagens are produced by fibroblasts in the interstitial matrix, and are most classically associated with fibrosis.^32^ For both C57BL/6 and BALB/c mice, expression of type I (*Col1a1*) and type III (*Col3a1*) collagen genes are most significant as compared to strain-specific controls. Furthermore, expression levels of these collagens in chemically induced BALB/c mice are significantly greater than those in chemically induced C57BL/6 mice (p=0.0108 and p=0.0386 for *Col1a1* and *Col3a1*, respectively). Numerous other collagens (e.g., 4a1, 5a3, and 6a3) were significantly induced by quinacrine treatment across one or both strains, with others trending upwards, as well. Interestingly, type I, III, and IV are amongst the collagens most clinically associated with severe fibrotic disease in the lung, liver, and kidney.^32^ These data show evidence of a greater fibrotic response in quinacrine-treated BALB/c mice compared to the other induction methods and compared to responses in C57BL/6 mice, also suggesting BALB/c mice are more susceptible to intrauterine fibrosis.

### 2.4. Greater induction of immune responses observed for fibrosis-induced BALB/c mice

Relative expression of a variety of immune relevant genes was also quantified, with p-values for all presented immune genes shown in **Supplemental Table 3**. Comparing induction methods to their strain-specific control group, chemically induced BALB/c mice showed the most highly induced immune cell gene expression levels (**Fig. 5A**). This was especially true for macrophage, B cell, and T cell markers. Further, these immune cell genes showed elevated expression for BALB/c curettage-induced mice compared to BALB/c control mice; however, quinacrine treatments induced greater and more significant expression differences. Although the induction methods damage tissues in different ways (i.e., mechanical damage versus chemical toxicity), this suggests similar immune mechanisms may lead to fibrosis in uterine tissues. With that said, further assessment is required.

**Figure 5.**
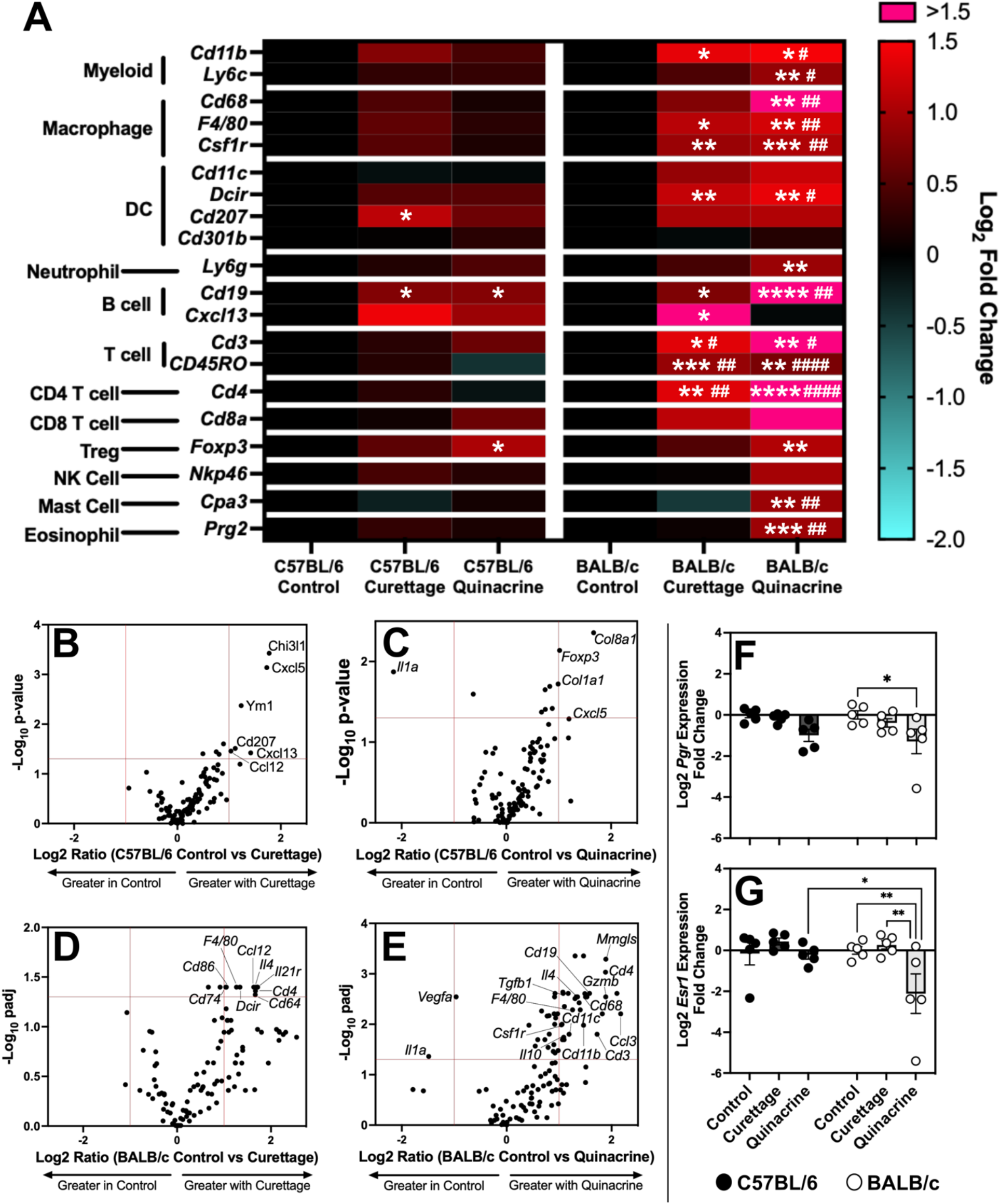
Quinacrine treatments in BALB/c mice stimulate the greatest expression of fibrotic and immune cell genes. Relative expression data for (**A**) immune cell-relevant genes quantified by NanoString are shown as average Log2 fold change values normalized to the strain specific control (DMPA treated/no induction) (n=4/group). Statistical significance was determined by ordinary two-way ANOVA with Tukey’s multiple comparisons test. Significant differences in gene expression compared to strain-specific control group are shown by *=p<0.05, **=p<0.01, ***=p<0.001, and ****=p<0.0001. Significant increases in gene expression compared across mouse strain for the same treatment are shown by ^#^=p<0.05, ^##^=p<0.01, ^###^=p<0.001, and ^####^=p<0.0001. Volcano plots are used to show significant (p>0.05) and differentially expressed (Log2 ratio>|2|) genes assessed by NanoString, comparing DMPA synchronized (**B**) C57BL/6 curettage to C57BL/6 control, (**C**) C57BL/6 quinacrine to C57BL/6 control, (**D**) BALB/c curettage to BALB/c control, and (**E**) BALB/c quinacrine to BALB/c control with notable genes labeled (averaged for n=4 mice/group). Students’ t-test used to determine p-values and adjusted p-values. Additional gene expression analysis by qPCR is shown for (**F**) Pgr and (**G**) Esr1 (n=5 mice/group, plotted average ± SEM). Statistical significance was determined by ordinary two-way ANOVA with Tukey’s multiple comparisons test, with *=p<0.05 and **=p<0.01.

Volcano plots illustrate genes with significant (p<0.05) and elevated magnitude expression (Log2>1) of all screened genes in the NanoString panel for each induction method (**Fig 5B-E**). Compared to C57BL/6 induction mice (**Fig. 5B&C**), more fibrosis and immune-relevant genes were most significantly upregulated in BALB/c curettage and chemically induced mice compared to their strain-specific control group (**Fig. 5D&E**). In order of magnitude of expression, *Ccl3, Col6a5, Mmgls, Gzmb, Cd4, Spp1, Cd3, Cd19, Il21r, Cd68, Col1a1, Fap, Cd11b, Ccl6, Dcir, Irf5, Il4, Stab1, Col3a1, F4/80, Cd11c, Pf4, Vegfd, Pgf, Il10, Tgfb1, Cd206, Tnfr2, Foxp3, and Csf1r* all show significantly increased expression in chemically induced uterine tissues as compared to control samples (**Fig. 5E**). Comparisons of induction method for BALB/c mice were plotted with adjusted, rather than standard, p-values as a more rigorous screening of significant gene responders.

Considering the possible effects of female sex hormone signaling on immune responses, qPCR was used to specifically measure relative gene expression of progesterone receptor (*Pgr*, **Fig. 5F**) and estrogen receptor (*Esr1*, **Fig. 5G**). Immune cells express sex hormone receptors;^33^ therefore, we hypothesized that elevated immune cell gene expression may correspond with elevated expression of *Pgr* and *Esr1*. However, quinacrine-treated BALB/c mice (again, with the greatest levels of fibrotic and immune cell gene expression) instead showed a significant decrease in both *Pgr* and *Esr1* expression as compared to control BALB/c mice (p=0.0186 and 0.0048 for *Pgr* and *Esr1*, respectively), and also when compared to quinacrine-treated C57BL/6 mice (p=0.0101 for *Esr1*). Female sex hormones, specifically estrogens, generally have a protective role against organ fibrosis, with ovariectomized *in vivo* models and post-menopausal women having greater susceptibility to fibrotic diseases affecting the kidney, liver, lung, and heart.^33^ Decreased receptor expression may decrease protective signaling in the highly fibrotic quinacrine-treated BALB/c uterine tissues, thereby increasing susceptibility. Although an interesting observation, future studies should investigate this potential pathway and confirm the observed decreased expression is conserved at the protein level, rather than an artifact of RNA dilution due to tissue infiltration by other responding cells.

Statistical analyses were also conducted to compare gene expression differences between strains for control, curettage, and chemically induced mice to assess factors which may contribute to fibrosis resistance or susceptibility (**Fig. 6**). Across all groups, C57BL/6 mice showed greater and more significant expression of the gene *resistin-like alpha* (*Retnla)* compared to BALB/c mice (p=0.0262 for control, p=0.0025 for curettage, and p=0.0027 for quinacrine, **Fig. 6A-C**). *Retnla is* induced during Th2-like immune responses in allergy and parasite infection, and has been described as a hallmark gene of alternative macrophage activation.^34,35^ The human genome does not specifically include *Retnla*; however, the resistin-like molecules family is conserved in mammals.^34^ In a study of helminth lung infections, greater fibrotic responses were measured for *Retnla*^-/-^ mice, suggesting a fibrosis modulatory function of *Retnla*.^35^ Such findings may suggest a potential role of *Retnla* in intrauterine fibrosis resistance in C57BL/6 mice, and could be a promising target for fibrosis treatment. However, further studies are needed to assess the tissue-specific effect of *Retnla* or similar clinically relevant genes.

**Figure 6.**
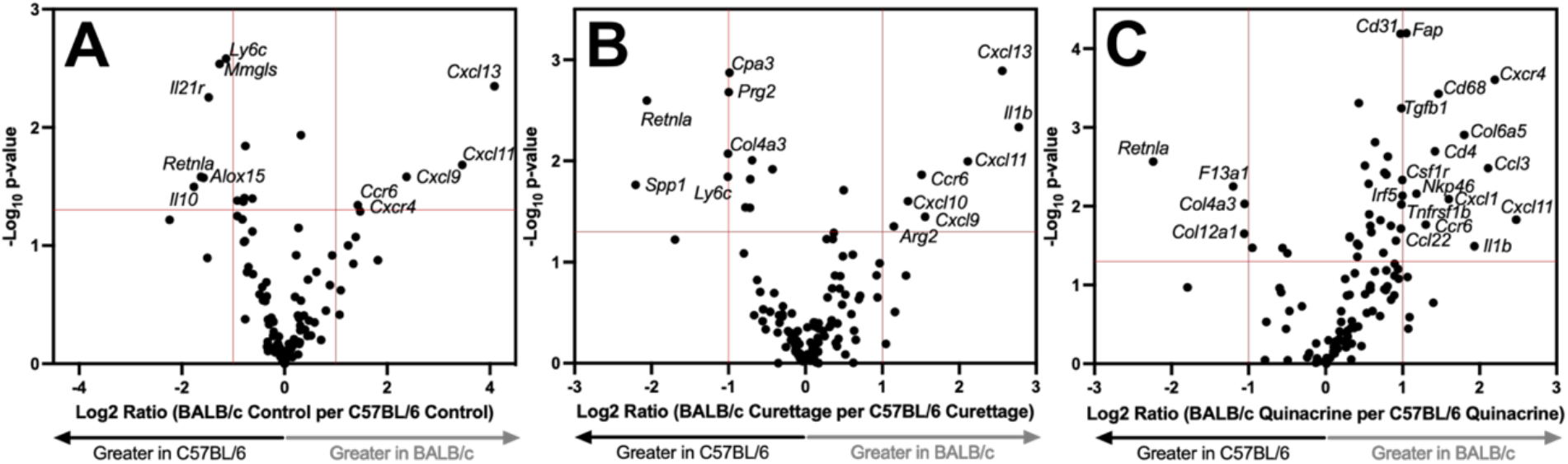
Macrophage-associated and pro-inflammatory genes are expressed greater in BALB/c than C57BL/6 mice. Volcano plots showing significant (p>0.05) and differently expressed genes (Log2 ratio>|2|) between DMPA controlled C57BL/6 and BALB/c mice for (**A**) control, (**B**) curettage, and (**C**) quinacrine treatments. Gene expression measured by NanoString, and p-values determined by t-test (n=4 mice per group).

In addition to immune cell genes previously discussed, quinacrine-induced BALB/c uterine tissues also showed elevated expression with significant or near significant differences in genes associated with inflammatory response, as compared to C57BL/6 mice (in order of expression magnitude: *Cxcl11, p=0.0148; Ccl3, p=0.0033; Cxcr4, p=0.0002; Il1b, p=0.0322; Cxcl1, p=0.0081; Tnfa, p=0.0538*; *Ccl22, p=0.0194; Cd31, p=0.00006;* **Fig. 6C**). Transforming growth factor-β (*Tgfb*) is typically considered an anti-inflammatory cytokine, but also has a significant role in fibrogenesis,^11^ and is also elevated in BALB/c (over C57BL/6) mice with chemical induction. Expression of these genes suggests a possible pro-inflammatory role of immune cells like macrophages in intrauterine fibrosis susceptibility in BALB/c mice. Interestingly, proposed mechanisms of quinacrine intrauterine fibrosis induction has been described as a uniquely human pathway, causing epithelial cell death, immune cell responses and release of cytokines like IL-1, TNF-α, TGF-β, and finally collagen deposition within the fallopian tube lumen.^7^ Although studies have stated that this response has not been replicated in rat, pig, and non-human primate models,^7,25^ similar immune factors and occlusion responses are observed here in BALB/c uterine tissues, supporting the suitability of this model.

### 2.5. Immune cell populations have the greatest frequency in BALB/c uterine tissues

Elevated presence of various immune cells was further assessed at the protein and single cell level by flow cytometry. Consistent with gene expression data, significantly greater leukocytes are detected in BALB/c tissues, with induction models having more leukocytes than control tissues, and chemically induced BALB/c tissues having the greatest leukocyte population (**Fig. 7A**). Of leukocytes, myeloid cell populations are similar across strains for control treatments (p=0.8293), and slightly but significantly elevated in BALB/c induction models (p=0.0462 for curettage and p=0.0057 for quinacrine, **Fig. 7B**). Of myeloid cells, we were especially interested in macrophage frequencies considering gene expression data and known dictating roles of macrophages in other fibrotic pathologies.^36^ F4/80 was used as a general marker of murine macrophages, and while CD68 is also sometimes cited as a pan-macrophage marker, other studies have used CD68 as a marker for more pro-inflammatory, M1/Th1-like macrophage phenotypes.^37,38^ While CD68^-^ macrophage populations are greater in C57BL/6 tissues (**Fig. 7C**), CD68^+^ macrophages have significantly greater populations in BALB/c tissues (**Fig. 7D**). T cell populations were also similar across strains and induction treatments, except for a significantly lower frequency in chemically induced BALB/c versus chemically induced C57BL/6 mice (p=0.0359, **Fig. 7E**). Despite this, CD4^+^ helper T cell populations had higher frequencies in BALB/c uterine tissues than C57BL/6 for all induction treatments (**Fig. 7F**). Both macrophages and CD4^+^ T cells are known to have a role in other fibrotic pathologies, and can synergistically propagate responses through cytokine signaling and activation by antigen presentation.^10,11^ Cell populations including natural killer cells, B cells, CD8^+^ T cells, and Tregs were also assessed; however, these populations did not have significant differences across treatment groups, or had low detected cell counts (**Supplemental Fig. 3**).

**Figure 7.**
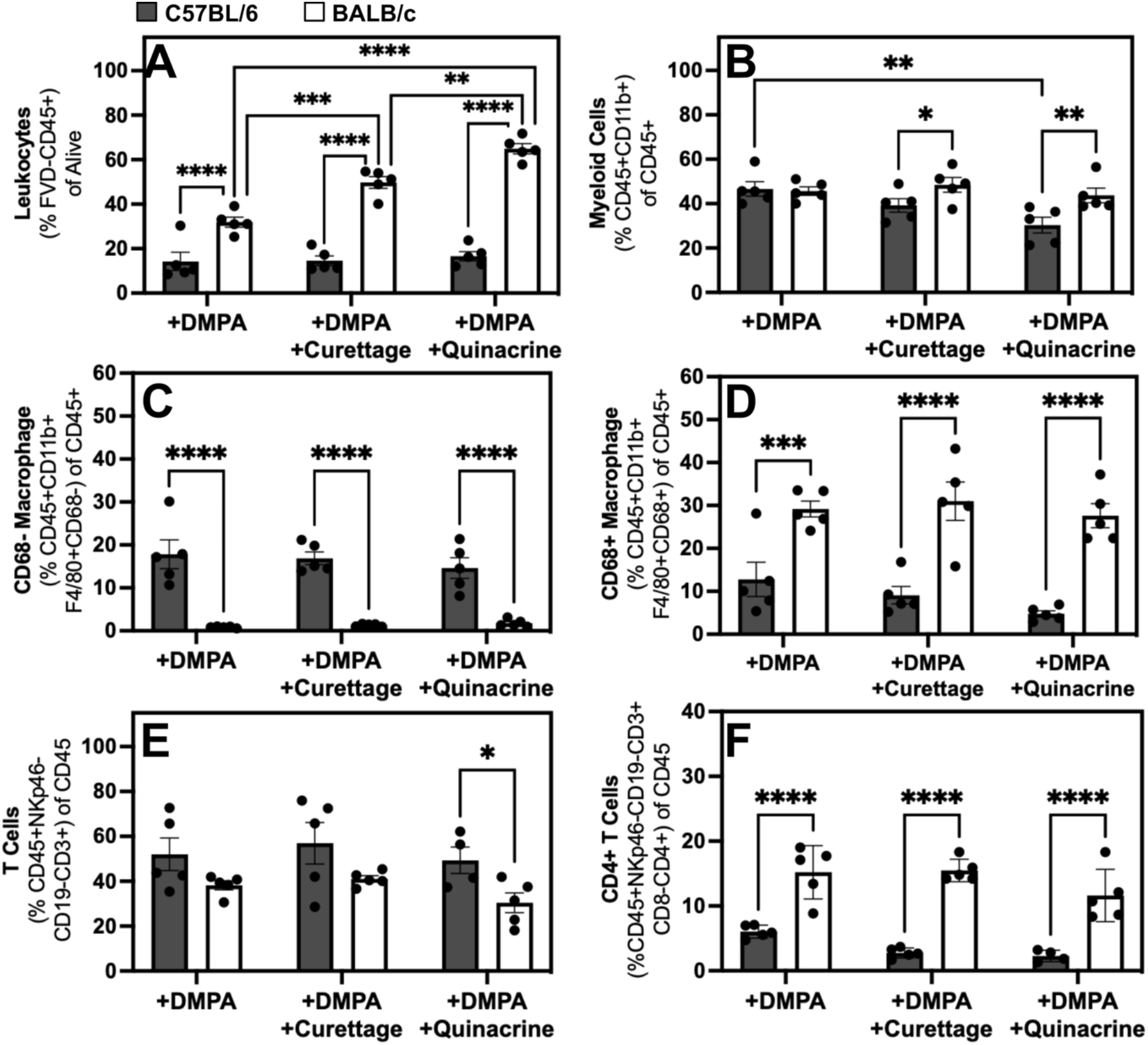
Greater populations of lymphocytes, pro-inflammatory CD68^+^ macrophages, and CD4^+^ T cells are found in BALB/c fibrosis induced uterine horn tissues. Uterine tissue immune cell populations for (**A**) leukocytes, (**B**) myeloid cells, (**C**) CD68^-^ macrophages, (**D**)CD68^+^ macrophages, (**E**) T cells, and (**F**) CD4^+^ T cells were measured by flow cytometry (n=5 mice, plotted average ± SEM). Statistically significant differences between induction treatment and between strains were determined by ordinary two-way ANOVA with Tukey’s multiple comparisons test, with *=p<0.05, ***=p<0.001, and ****=p<0.0001.

Gene expression and flow cytometry data suggest that pro-inflammatory macrophages may be a key immune population in intrauterine fibrosis susceptibility. Published studies of human tissues have proposed conflicting roles of macrophages in IUA development, however these works differ in their methods to mitigate the effects of the menstrual cycle. Liu *et al*. studied Asherman’s syndrome patient tissues collected at the late proliferating phase – when estrogen and progesterone concentrations are low – and found decreased macrophage populations in fibrotic tissues. Their findings suggested a critical role of macrophages in endometrial repair, rather than fibrosis provocation.^39^ Conversely, Santamaria *et al*. synchronized patient cycles using hormone replacement therapy prior to endometrial biopsy and found a highly significant increase in macrophages, and specifically pro-inflammatory macrophages, in Asherman’s syndrome tissues.^3^ While hormone replacement treatments may alter native immuno-endocrine dynamics, studies of the natural cycle may not be appropriate as moderate-to-severe Asherman’s syndrome patients typically do not undergo menstruation.^3^ Moreover, post-partum patients appear to have the greatest risk of IUA induction,^1^ when plasma progesterone and estrogen is high and beginning to decrease.^40^ Hormone treatments therefore may be a better model of a fibrosis induction-susceptible microenvironment, but specific mechanisms of hormone signaling in early tissue responses should be studied further. Elevated pro-inflammatory macrophage populations in our DMPA-synchronized, chemically induced BALB/c intrauterine fibrosis model therefore appears consistent with human studies, and could be a more promising model of clinical response than C57BL/6 mice with similar treatments.

## 3. Conclusions

In this study, we compared multiple strains and induction methods of intrauterine fibrosis to determine a consistent and more clinically relevant mouse model to study IUA. Further, DMPA synchronization was found to improve response consistency. With DMPA administration, the chemically induced mouse model established here proved to be a simple, well-tolerated, and effective method for intrauterine fibrosis induction. Moreover, BALB/c mice showed greater intrauterine fibrosis susceptibility than C57BL/6 mice. In these highly fibrotic uterine tissues, greater CD68^+^ macrophage and CD4^+^ T cell responses were identified via gene expression analyses and flow cytometry. Consistently greater expression of the fibrosis suppressive gene *Retnla* was also identified in C57BL/6 mice compared to BALB/c, suggesting a possible fibrosis resistance mechanism. As such, the DMPA-synchronized, chemically induced BALB/c intrauterine fibrosis model reported here proved to not only be robust and reliably effective, but also possibly more clinically faithful, showing promise for improving our understanding of IUA as well as for use in preclinical assessment of potential therapies.

## 4. Methods

### 4.1. Animal use and care

Procedures and mouse studies were approved by the Johns Hopkins Animal Care and Use Committee (ACUC). Mice were 8-week-old, female, C57BL/6 or BALB/c mice (Jackson Laboratory). For hormone synchronized studies, mice underwent subcutaneous injections of 3 mg depot medroxyprogesterone (DMPA, 150mg/mL Amphastar Pharmaceuticals, Cat# 0548-5400-00), administered every four weeks. Mice were sacrificed 28 days after initial induction.

### 4.2. Transcervical administration of intrauterine sclerosing agents

In initial pilot studies, silver nitrate (Chem-Impex, Cat# 272390247) was delivered in sterile cell culture grade water (HyPure, HyClone Laboratories, Cat# SH30538.02) to prevent precipitate formation with sodium chloride. Quinacrine hydrochloride hydrate (Cayman Chemical, Cat# 15041) was suspended in sterile saline (ICU Medical, Cat# 98964). All solutions were transcervically administered at a 1.6 mL/kg dose. DMPA-synchronized quinacrine treatment studies used the 350 mg/kg dose, administered initially and after 14 days. Mice were anesthetized using isoflurane, and the quinacrine solution was transcervically administered using a cryopreservation and insemination device for mice (mC&I, ParaTechs, Cat# 60020) following device instructions.

### 4.3. Surgical curettage of uterine horn

Mice were anesthetized using isoflurane and given a subcutaneous injection of 5 mL/kg lactated ringers (AmeriSource Bergen, Cat# 795309) and buprenorphine (Wedgewood Pharmacy) as an analgesic. Aseptic technique was followed. A single skin incision was made just off the midline of the dorsum, approximately half-way between the bottom of the ribcage and base of the tail. Following blunt dissection, a small incision was made in the muscle layer above the right ovarian fat pad. The right uterine horn was then pulled through the incision, and curettage conducted via 10x insertion and rotation of a 25G needle (BD Medical, Cat# 305122) into the horn lumen. After curettage, the uterine horn was returned through the incision, the incision was sutured. The curettage procedure was then repeated for the left uterine horn. The skin incision was closed with surgical staples and wound glue.

### 4.4. Uterine tissue images and histology

Intact, resected female reproductive tracts were collected from the mice at sacrifice (with fat tissue removed), and the gross anatomy of the tissues captured via photography with a ruler for scale. Uterine horn cross sections (two per horn per mouse) were excised and fixed in a 4% (v/v) formaldehyde solution (Electron Microscopy Sciences, 32% aqueous solution, Cat# 100496-496) in PBS. Tissues were processed, sectioned, and stained with Masson’s trichrome and hematoxylin and eosin (H&E) by the Johns Hopkins Oncology Tissue Services core. Microscopy images were collected using brightfield microcopy using a ThermoFisher EVOS XL Core Imaging System.

### 4.5. Gene expression analyses

Uterine tissue sections used for gene expression analyses were liquid nitrogen snap-frozen immediately following dissection. RNA was isolated from these tissue samples using TRIzol (Invitrogen, Cat# 15596018) according to manufacturer instructions and using a homogenizer (Polytron PT 2500E with Polytron 12 mm Generator, Kinematica, Inc). Total sample RNA concentrations were measured using a NanoDrop 2000 spectrophotometer (Thermo Scientific).

For quantitative polymerase chain reaction (qPCR) total RNA was diluted to a concentration of 1 µg RNA/12.2 µL total sample volume in DEPC-water (Invitrogen, Cat# 750024). All RNA samples first underwent a DNAse step (1 µL DNAse/12.2 µLRNA, RQ1 RNAse-Free DNAse, Promega), then reverse transcription using a High-Capacity cDNA Reverse Transcription kit (Thermo Fisher Scientific, Cat# 4368814) according to manufacturer instructions using a SimpliAmp Thermal Cycler (Thermo Fisher Scientific). Resulting cDNA was mixed with 8 µL Power SYBR Green PCR Master Mix (Thermo Fisher Scientific, Cat# 4368702), 1 µL forward and reverse primers (**Table 1**), and 1.2 µL Milli-Q water, and amplified with three technical replicates at a 5 µL/well volume within a 384-well plate. Using a QuantStudio5 real-time PCR system (Thermo Fisher Scientific), samples were incubated at 95°C for 10 min, then underwent 40 cycles of heating at 95°C for 15 seconds then 60°C for 1 minute. The comparative ddCT method was used to analyze qPCR data, using β-actin (*Actb*) as the housekeeping gene and control tissue groups (specified in figure legends) for normalization.

For NanoString analysis, a custom-designed, 120-gene panel was used to identify RNA expression related to various aspects of immune responses. Total RNA per sample was diluted to 5 µg RNA/50 µL DEPC-water, and RNA samples were processed according to manufacturer instructions. Absolute copy number of RNA for the genes of interest were quantified using an nCounter (NanoString Technologies).

### 4.6. Uterine tissue digestion, cell staining, and flow cytometry

Sections of each uterine horn were added to 1 mL of RPMI 1640 media (Gibco, Cat# 61870127) during collection. Digestion methods were adapted from Depierreux, *et al*.^41^, using 3 mL of pre-warmed (37°C) digestion media consisting of RPMI with 0.52 WU/mL Liberase Thermolysin Medium (Roche, Cat# 5401020001) and 30 µg/mL DNAse I (grade II from bovine pancreas, Roche, Cat# 10104159001) added to each sample tube. Samples were incubated in a water bath at 37°C for 30 minutes, vortexing every 5 minutes to mix. Samples then underwent mechanical dissociation using “program C” on a gentle MACS dissociator (Miltenyi Biotech) and were passed through a 40 µm filter with 2 mL Dulbecco’s phosphate-buffered saline (PBS, Gibco, Cat# 14190144) added. Cell suspensions were centrifuged at 400 g for 10 min at 4°C, and pellets underwent red blood cell lysis (eBioscience 1x RBC Lysis Buffer, Invitrogen, Cat# 00-4333-57) according to manufacturer instructions. Solutions were centrifuged at 400g for 10 min at 4°C, supernatant aspirated, the pellet was resuspended in 1 mL flow cytometry staining buffer (Invitrogen, Cat# 00-4222-26), and cells were counted.

Cells were plated at approximately 0.5×10^6^ per well in a 96-well plate for all individually stained samples and as pooled cell samples for fluorescence minus one (FMO) controls. Plates were centrifuged at 400g for 10 min at 4°C, supernatant removed, 50 µL of Fc block solution (20 µL/mL CD16/CD32 Monoclonal Antibody (Invitrogen, Cat# 14-0161-86) in stain buffer) was added to each well, and incubated on ice for 10 minutes. Two separate cell plates were used for innate and adaptive immune cell panels. Optimal antibody concentrations were determined via titrations. In both panels, fixable viability dye (FVD, eFluor 780, Invitrogen, Cat# 65-0865-14, 1:400 dilution) was used to stain for alive cells, and a fluorescently conjugated anti-mouse antibody against CD45 (PerCP, BioLegend, clone# 30-F11, Cat# 103130, 1:400 dilution) for lymphocytes. Myeloid panel specific anti-mouse antibodies included CD11b (Alexa Fluor 488, BioLegend, Clone# M1/70, Cat# 101217, 1:3200 dilution), CD11c (Brilliant Violet 421, BioLegend, Clone# N418, Cat# 117330, 1:25 dilution), CD68 (Alexa Fluor 647, BioLegend, Clone# FA-11, Cat# 137004, dilution 1:400), F4/80 (Brilliant Violet 510, BioLegend, Clone# BM8, Cat# 123135, 1:25 dilution), Ly6g (Alexa Fluor 700, BioLegend, Clone# 1A8, Cat# 127622, 1:400 dilution), SiglecF (Super Bright 600, Invitrogen, Clone# 1RNM44N, Cat# 63-1702-82, 1:400 dilution), and FceR1 (PE, Invitrogen, Clone# MAR-1, Cat# 12-5898-83, 1:100 dilution). Lymphocyte panel specific antibodies included NKp46 (Brilliant Violet 650, BioLegend, Clone# 29A1.4, Cat# 137635, 1:25 dilution), CD19 (Brilliant Violet 510, BioLegend, Clone# 6D5, Cat# 115546, 1:25 dilution), CD3 (Alexa Fluor 488, BioLegend, Clone# 17A2, Cat# 100210, 1:25 dilution), CD4 (Alexa Fluor 700, BioLegend, Clone# GK1.5, Cat# 100430, 1:200 dilution), CD8a (Brilliant Violet 711, BioLegend, Clone# 53-6.7, Cat# 100759, 1:50 dilution), FoxP3 (Alexa Fluor 647, BioLegend, Clone# 150D, Cat# 320014, 1:12.5 dilution), and CD25 (PE, BioLegend, Clone# 3C7, Cat# 101904, 1:25 dilution). 50 µL of surface labeling antibody solutions were added at appropriate concentrations in staining buffer with 100 µL/mL Super Bright Complete Staining Buffer (Invitrogen, Cat# SB-4401-75) and incubated for 30 minutes. Concurrently, single stained beads (UltraComp eBeads Plus Compensation Beads, Invitrogen, Cat# 01-3333-42) were stained with 1 µL antibody per 100 µL staining buffer, and ArC Amine Reactive Compensation Beads (Invitrogen, Cat# A10346) were stained with 1 µL FVD per 100 µL PBS according to manufacturer specifications. Cyto-Fast Fix/Perm Buffer Set (BioLegend, Cat# 426803) and Cyto-Last buffer (BioLegend, Cat# 422501) were used according to manufacturer instructions to fix and permeabilize cells in the myeloid panel. Foxp3/Transcription Factor Staining Buffer Set (Invitrogen, Cat# 00-5521-00) was used according to manufacturer instructions for the lymphocyte panel. The following day, cells were stained with intracellular antibodies (CD68 and FoxP3) and incubated for 30 min. Prior to measurement, cells were washed and resuspended in 200 µL staining buffer. Cells were measured on a Attune NxT Flow Cytometer (ThermoFisher). Data were analyzed using FlowJo (version 10.10.0) software.

### 4.7. Statistical Analysis

ROSALIND, Inc. NanoString Gene Expression analyses was used to analyze NanoString data, including t-tests to assess induction method versus control, and strain comparisons of relative expression for all screened genes. All other analyses were conducted using Prism (version 10.2.3, GraphPad). Specific statistical tests and replicate numbers are detailed in the figure legends.

## Data Availability

The datasets generated during and/or analyzed during the current study are available from the corresponding author on reasonable request.

## Acknowledgements

This work was funded with support from the Ovarian Cancer Research Alliance (OCRA) (Grant #ECIG-2024-3-1556) as well as with support from the Biomedical Engineering Department at Johns Hopkins University. J.L.H. and J.C.D. were supported by the Academic Success via Postdoctoral Independence in Research and Education (ASPIRE)/ National Institutes of Health (NIH) Institutional Research and Academic Career Development Award (IRACDA) program (NIH K12-GM123914). J.L.S. was supported by a fellowship from the National Science Foundation (DGE-1746891). J.L.S. and V.V.S. were both supported by NIH F31 graduate fellowships, and N.R.P. was supported by a Johns Hopkins Provost’s Undergraduate Research (PURA) award. The authors would like to also acknowledge the Johns Hopkins Sidney Kimmel Comprehensive Cancer Oncology Tissue Services for processing the histology used in this manuscript as well as the Johns Hopkins Research Animal Resources for animal care and support for this work.

## Authorship Contributions

J.L.H. conceptualized the study including methodology, conducted the formal analysis, and wrote the original manuscript draft including figure visualizations. J.D. contributed to experimental procedures, conducted and contributed to the analysis of initial gene expression data. J.L.S. contributed to validation and analysis of flow cytometry data. N.R.P. collected histology images. V.V.S. contributed to experimental procedures and design of qPCR primers. J.C.D. contributed to the conceptualization of the study, analysis, project administration, provided resources and funding acquisition. All authors reviewed and edited the manuscript draft.

