## Supplementary Information for "Immunologic comparisons of strain and induction method in an improved mouse model of intrauterine fibrosis"

***Supplemental Table 1. p-values determined for mouse % weight change***

| | | | |
|---|---|---|---|
| | **C57BL/6 mice** | | |
| **Time (d)** | **Curettage vs Control** | **Quinacrine vs Control** | **Curettage vs Quinacrine** |
| 1 | ** (0.004) | ** (0.0043) | NS (0.6915) |
| 2 | ** (0.0093) | ** (0.0083) | * (0.0426) |
| 3 | * (0.0181) | ** (0.0058) | NS (0.0939) |
| 14 | * (0.0122) | NS (0.228) | NS (0.9994) |
| 21 | NS (0.3411) | NS (0.8694) | NS (0.8389) |
| 28 | NS (0.6183) | NS (0.4986) | NS (0.9668) |
| | **BALB/c mice** | | |
| **Time (d)** | **Curettage vs Control** | **Quinacrine vs Control** | **Curettage vs Quinacrine** |
| 1 | NS (0.0649) | NS (0.0529) | NS (0.4872) |
| 2 | * (0.022) | NS (0.8122) | NS (0.4727) |
| 3 | * (0.0434) | NS (0.0710) | NS (0.4232) |
| 14 | NS (0.0574) | NS (0.38) | NS (0.2101) |
| 21 | NS (0.4302) | NS (0.3048) | NS (0.8358) |
| 28 | NS (0.2511) | NS (0.3266) | NS (0.8601) |

*Determined by repeated measures two-way ANOVA with Geisser-Greenhouse correction and Tukey's multiple comparisons test. For p>0.05 = NS (non-significant), p<0.05 = *, and p<0.01 = **.*

**Supplemental Table 2. p-values (2-way ANOVA) of all collagen genes in heat map**

| Gene | p-value for method comparison against strain-specific control | | | | p-value for strain comparison | |
|---|---|---|---|---|---|---|
| | C57BL/6 Curettage vs Control | C57BL/6 Quinacrine vs Control | BALB/c Curettage vs Control | BALB/c Quinacrine vs Control | Curettage C57BL/6 vs BALB/c | Quinacrine C57BL/6 vs BALB/c |
| *Col1a1* | NS (0.9998) | **** (<0.0001) | NS (0.9973) | **** (<0.0001) | NS (0.9293) | * (0.0108) |
| *Col2a1* | NS (0.8927) | NS (0.8817) | NS (0.7412) | NS (0.1770) | NS (0.7763) | NS (0.1822) |
| *Col3a1* | NS (0.8987) | *** (0.0006) | NS (0.5468) | **** (<0.0001) | NS (0.5404) | * (0.0386) |
| *Col4a1* | NS (0.4859) | NS (0.1590) | NS (0.7187) | *** (0.0003) | NS (0.7029) | ** (0.008) |
| *Col4a3* | NS (0.0935) | NS (0.9104) | NS (0.0642) | NS (0.99950) | NS (0.8453) | NS (0.7055) |
| *Col5a2* | NS (0.0935) | NS (0.9104) | NS (0.0642) | NS (0.9995) | NS (0.8453) | NS (0.7055) |
| *Col5a3* | NS (0.6485) | * (0.0110) | ** (0.0053) | *** (0.0002) | * (0.0141) | NS (0.0723) |
| *Col6a1* | NS (0.9346) | NS (0.2069) | NS (0.1568) | NS (0.0742) | NS (0.1298) | NS (0.5713) |
| *Col6a2* | NS (0.3363) | NS (0.4927) | NS (0.0475) | NS (0.1528) | NS (0.2750) | NS (0.4364) |
| *Col6a3* | NS (0.8586) | * (0.0306) | NS (0.2124) | NS (0.0630) | NS (0.2353) | NS (0.7224) |
| *Col6a5* | NS (0.7275) | NS (0.9975) | NS (0.0591) | NS (0.0518) | NS (0.1062) | * (0.0239) |
| *Col8a1* | NS (0.2728) | NS (0.0142) | NS (0.8331) | NS (0.0267) | NS (0.3224) | NS (0.7674) |
| *Col12a1* | NS (0.6796) | NS (0.3754) | NS (0.9826) | NS (0.7281) | NS (0.3189) | NS (0.5515) |
| *Col24a1* | NS (0.9999) | NS (0.9641) | NS (0.7257) | NS (0.3846) | NS (0.4429) | NS (0.1242) |

*Determined by ordinary two-way ANOVA with Tukey's multiple comparisons test. For p>0.05 = NS (non-significant), p<0.05 = *, p<0.01 = **, p<0.001 = ***, and p<0.0001 = ****.*

**Supplemental Table 3. p-values (2-way ANOVA) of all immune cell genes in heat map**

| Gene | p-value for method comparison against strain-specific control | | | | p-value for strain comparison | |
|---|---|---|---|---|---|---|
| | C57BL/6 Curettage vs Control | C57BL/6 Quinacrine vs Control | BALB/c Curettage vs Control | BALB/c Quinacrine vs Control | Curettage C57BL/6 vs BALB/c | Quinacrine C57BL/6 vs BALB/c |
| *Cd11b* | NS (0.2472) | NS (0.6612) | * (0.0249) | * (0.0165) | NS (0.2326) | * (0.0398) |
| *Ly6c* | NS (0.5493) | NS (0.4936) | NS (0.2335) | ** (0.0097) | NS (0.5331) | * (0.0424) |
| *Cd68* | NS (0.5307) | NS (0.9445) | NS (0.1892) | ** (0.0061) | NS (0.4721) | ** (0.0046) |
| *F4/80* | NS (0.2588) | NS (0.7629) | * (0.0137) | ** (0.0051) | NS (0.1388) | ** (0.0089) |
| *Csf1r* | NS (0.0981) | NS (0.7628) | ** (0.0027) | *** (0.0008) | NS (0.1008) | ** (0.0014) |
| *Cd11c* | NS (0.9818) | NS (0.9945) | NS (0.3688) | NS (0.1785) | NS (0.1342) | NS (0.0653) |
| *Dcir* | NS (0.3145) | NS (0.2915) | ** (0.0072) | ** (0.0017) | NS (0.0634) | * (0.0189) |
| *Cd207* | * (0.0464) | NS (0.3176) | NS (0.0776) | NS (0.0626) | NS (0.7947) | NA (0.3574) |
| *Cd301b* | NS (0.9971) | NS (0.7280) | NS (0.9761) | NS (0.7834) | NS (0.7807) | NS (0.9247) |
| *Ly6g* | NS (0.7176) | NS (0.1807) | NS (0.2555) | ** (0.0048) | NS (0.4035) | NS (0.0872) |
| *Cd19* | * (0.0295) | * (0.0275) | * (0.0354) | **** (<0.0001) | NS (0.9293) | ** (0.0098) |
| *Cxcl13* | NS (0.0625) | NS (0.2764) | * (0.0341) | NS (0.9941) | NS (0.7648) | NS (0.1081) |
| *Cd3* | NS (0.8777) | NS (0.4121) | * (0.0327) | ** (0.0062) | * (0.0354) | * (0.0378) |
| *Cd45ro* | NS (0.5338) | NS (0.1411) | *** (0.0007) | ** (0.0042) | ** (0.0027) | **** (<0.0001) |
| *Cd4* | NS (0.7212) | NS (0.8793) | ** (0.0011) | **** (<0.0001) | ** (0.0022) | **** (<0.0001) |
| *Cd8a* | NS (0.9921) | NS (0.6959) | NS (0.3487) | NS (0.1590) | NS (0.2077) | NS (0.2814) |
| *Foxp3* | NS (0.1345) | ** (0.0039) | NS (0.2022) | ** (0.0029) | NS (0.8133) | NS (0.8894) |
| *Nkp46* | NS (0.6869) | NS (0.9024) | NS (0.9980) | NS (0.1487) | NS (0.4487) | NS (0.1418) |
| *Cpa3* | NS (0.6119) | NS (0.8960) | NS (0.2335) | ** (0.0064) | NS (0.4694) | ** (0.0064) |
| *Prg2* | NS (0.3367) | NS (0.7484) | NS (0.9443) | *** (0.0007) | NS (0.2737) | ** (0.0012) |

*Determined by ordinary two-way ANOVA with Tukey's multiple comparisons test. For p>0.05 = NS (non-significant), p<0.05 = *, p<0.01 = **, p<0.001 = ***, and p<0.0001 = ****.*

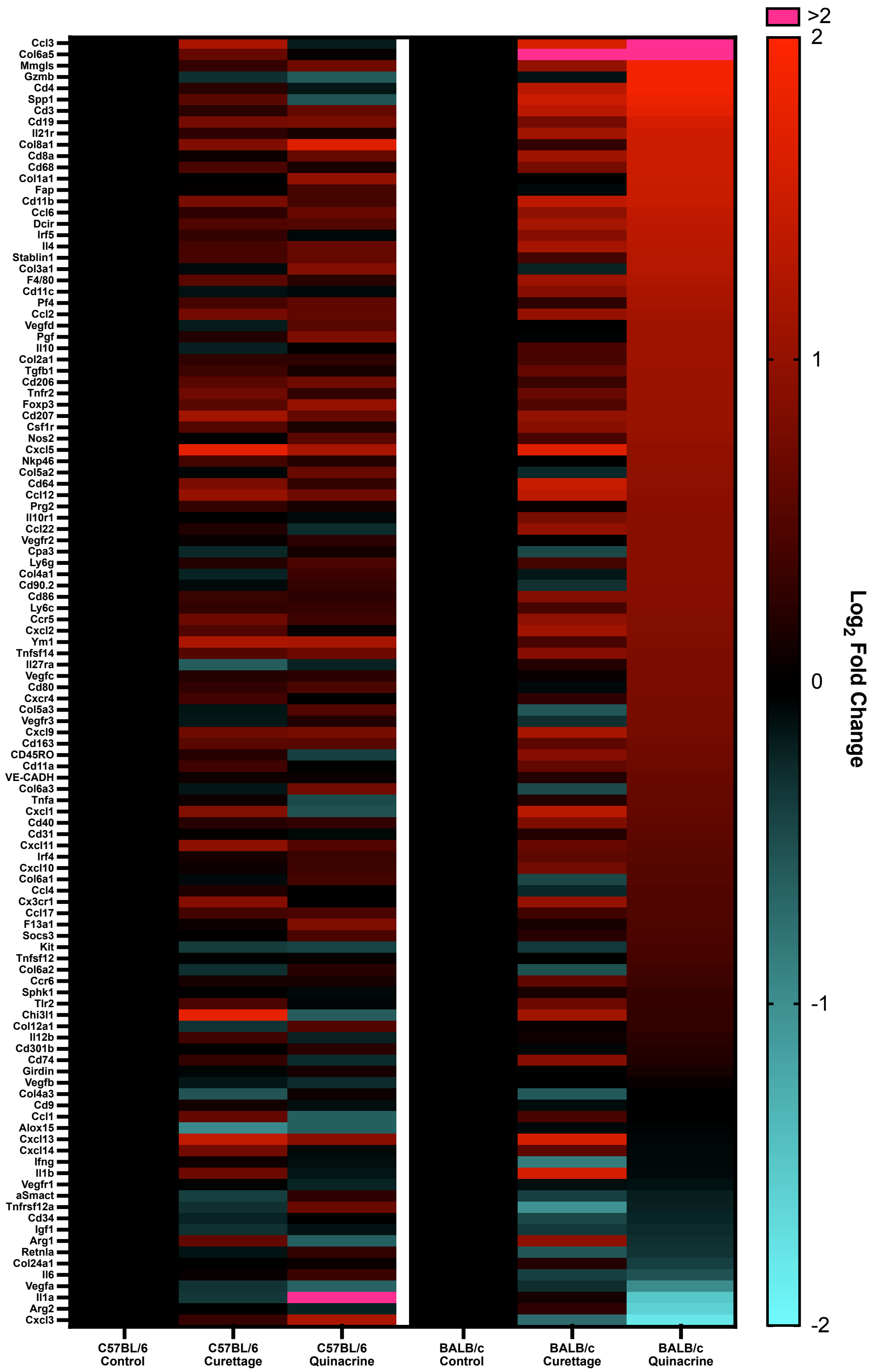


**Supplemental Figure 1. Heat map of all NanoString-measured gene expression.** Heat map of average $Log_2$ fold change values of all measured genes normalized to strain specific control (n=4/group). Portions of this data correspond to the heatmaps shown in Figs. 4G & 5A.

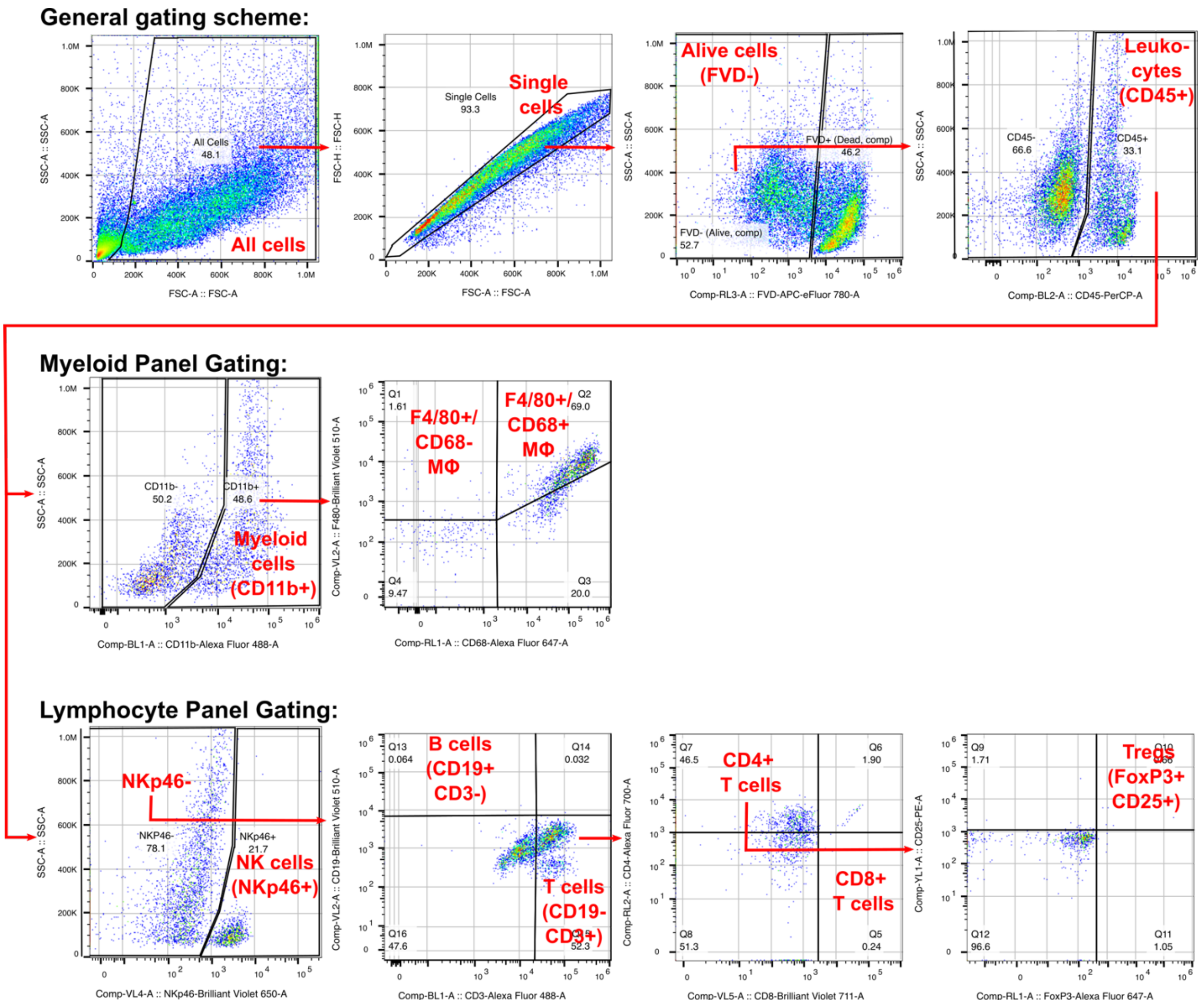


**Supplemental Figure 2. Flow cytometry gating strategy.** For both panels, cells were first gated away from debris and then gated as singlets. Negative staining for fixable viability dye (FVD) was used to identify alive cells. Of alive cells, positive staining for CD45 was used as a marker for leukocytes for sub-gating in both panels. For the myeloid panel, positive staining for CD11b was used to identify myeloid cells, and of CD11b$^+$ cells, F4/80 and CD68 were used as markers for mouse macrophages and pro-inflammatory-like macrophages, respectively. For the lymphocyte panel, natural killer cells were determined by positive staining of NKp46. B cells and T cells were sub-gated from NKp46$^-$ cells, using CD19 and CD3 as markers, respectively. Of CD3$^+$ T cells, CD4 and CD8 were used to further classify T-cell subtype. Tregs were classified by positive staining of FoxP3 and CD25 within CD4$^+$ T cells.

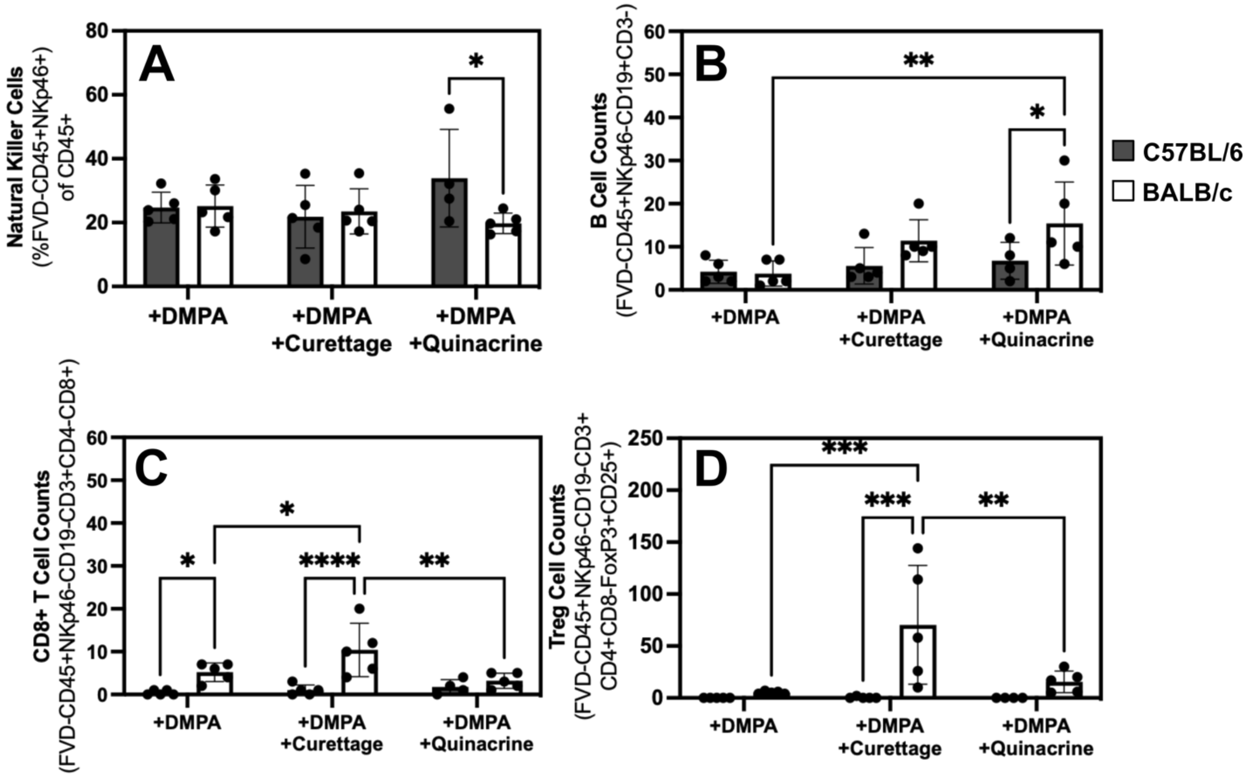


**Supplemental Figure 3. Additional lymphocyte cell populations are measured.** Flow cytometry data for (**A**) % Natural killer cells (% of CD45$^+$ cells), (**B**) B cell counts, (**C**) CD8$^+$ T cell counts, and (**D**) Treg counts. Statistically significant differences were determined by ordinary two-way ANOVA with Tukey's multiple comparisons test, with *=$p<0.05$, **=$p<0.01$, ***=$p<0.001$, and ****=$p<0.0001$ (plotted average ± standard deviation to compare sample group variation).
